# The Effect of Aldehyde Dehydrogenase Activator, Alda-1^®^, on the Ethanol-induced Brain Damage in a Rat of Binge Ethanol Intoxication

**DOI:** 10.1101/353854

**Authors:** Changsun Kim, Sejin Hwang, Hyuk Joong Choi, Tae Ho Lim, Ju-seop Kang

## Abstract

**Aims:** This study aimed to investigate whether an aldehyde dehydrogenase (ALDH) activator (Alda-1^®^) reduces neuronal damage in a rat model of binge ethanol exposure.

**Methods:** Thirty-six adolescent male rats (130-150 g) were randomly assigned into three groups: sham, ethanol-only group (25% ethanol intragastrically thrice daily for four days, approximately 10 g/kg/day) and ethanol with Alda-1^®^ group (10 mg/kg thrice daily for four days). The ALDH activity at baseline and 90 min after the last infusion in each group was measured. Brain damage was investigated using Luxol fast blue-Cresyl violet staining in the hippocampus, CA1 and CA2/3. The activation of astrocytes and microglia was examined using immunohistochemistry for antiglial fibrillary acidic protein (GFAP) and anti-ionized calcium-binding adapter molecule 1 (Iba-1).

**Results:** After a four-day binge, the ALDH activity level was doubled in the ethanol with Alda-1^®^ group (mean: 7.87, SD: 0.67), whereas the levels in the sham group (mean: 4.07, SD: 0.53) and ethanol-only group (mean: 3.77, SD: 0.36) were slightly decreased. More significant neuronal shrinkage, fewer neurons, and loss of Nissl in the hippocampus were observed in the ethanol-only group compared to the ethanol with Alda-1^®^ group. Astrocytosis and microgliosis of the hippocampus also showed increased activation in the ethanol only group compared with the ethanol with Alda-1^®^ group.

**Conclusion:** Alda-1^®^ administration reduces cytotoxic damage to the hippocampus in adolescent rats with binge ethanol exposure.

## Introduction

Excessive ethanol consumption affects many organ systems, including the central nervous system (CNS) ^1, 2^. It modifies brain function through its effects on brain tissue, neurotransmitter systems, and metabolism ^3^. Several studies have suggested that a binge-drinking pattern may play a central role in neuropathology ^4–6^, which is strongly associated with alcoholism or alcohol dependence ^7^.

Ethanol-induced cytotoxic damage can be attributed to acetaldehyde accumulation. Ingested ethanol is metabolized to acetaldehyde by alcohol dehydrogenase (ADH) (Figure 1). Acetaldehyde, a cytotoxic product of ethanol metabolism, produces reactive oxygen species and also reacts with proteins to form adducts ^8^. These adducts can further interact with other proteins and fatty acids to cause adverse metabolic effects ^9^.

**Figure 1.**
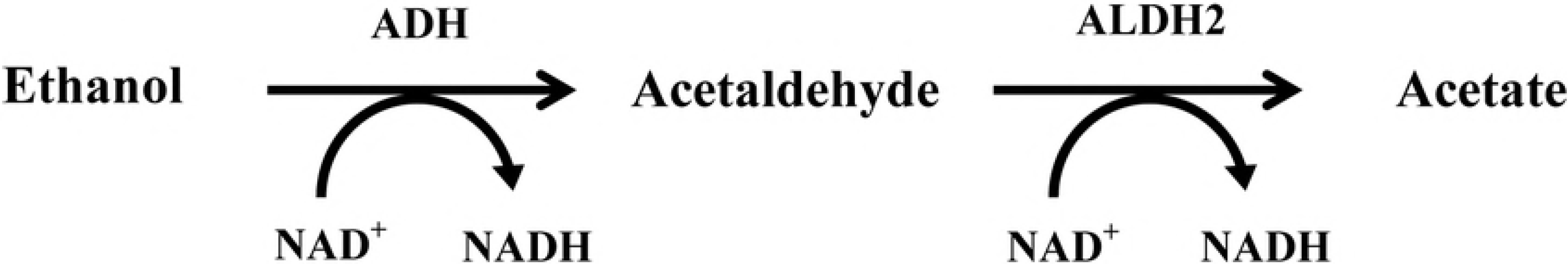
Ethanol metabolism: Ethanol is oxidized to acetaldehyde by the ADH-containing coenzyme NAD^+^, and acetaldehyde is further oxidized to acetate by ALDH2. NAD^+^ is converted to NADH during this process.

Acetaldehyde can be catalyzed to acetate by the mitochondrial aldehyde dehydrogenase 2 (ALDH2) enzyme. The deficiency of this enzyme leads to increased blood levels of acetaldehyde, which may increase the toxic effect of acetaldehyde. Numerous studies have demonstrated that blood acetaldehyde concentrations are significantly increased even after a moderate intake of ethanol in ALDH2-deficient subjects ^10–14^, and these subjects are susceptible to alcohol-induced CNS pathologies ^15^.

The accumulation of acetaldehyde plays an important role in cytotoxic oxidative injury. Thus, ALDH2-mediated detoxification of acetaldehyde is important to protect against acetaldehyde-induced oxidative injury. Several attempts have been made to ameliorate cytotoxic injury by increasing ALDH2 activity. Fu et al.^16^ suggested that increasing ALDH2 activity reduces cerebral ischemia/reperfusion injury in rats. Chen et al.^17^ demonstrated that activation of ALDH2 reduces ischemic damage to the heart and also reported the neuroprotective effect of ALDH2 activation in Parkinsonism ^18^. However, the ALDH2 activation-mediated neuroprotective effect of binge ethanol-induced brain injury has not been assessed.

We hypothesized that blood acetaldehyde levels could be reduced if ALDH activity is increased and that the neurocytotoxic effect of acetaldehyde could be ameliorated. We aimed to investigate whether an aldehyde dehydrogenase activator, Alda-1^®^, improves ethanol-induced neuronal damage in a rat model of binge ethanol exposure.

## Materials and Methods

### Animals

Thirty-six juvenile and adolescent male Wistar albino rats (postnatal day 30, weight: 130 to 150 g) were used in this study. The rats were maintained at 28°C with a 12-hour light-dark cycle and 5% humidity. The rats were permitted tap water at all times before and during experimental procedures. This study was approved by the Institutional Animal Care and Use Committee.

### Alda-1^®^ ALDH activator

Alda-1^®^ (ALDH activator-1, Sigma-Aldrich, USA) was used as an ALDH activator in this study. Alda-1^®^ is an ALDH2 agonist that stimulates acetaldehyde oxidation by ALDH2 by improving nicotinamide adenine dinucleotide (NAD) binding. Chen et al. ^17^ demonstrated that Alda-1 increased ALDH activity in subjects with wild type ALDH2 deficiency (ALDH2*1/*1 homotetramers) and Asian variant ALDH2 deficiency (ALDH2*2) by approximately 2.1-fold and 11-fold, respectively.

### Adolescent binge alcohol model and study design

A four-day binge model was first described by Majchrowicz in 1975 ^8^, and has been modified in several reports^4, 9, 19–21^. The binge rats received 25% ethanol with a nutritionally complete liquid diet (Ensure Plus^®^), and control rats were provided the same diet plus a calorie-matched dextrose solution via intragastric gavage every 8 hours for four days. Thirty-six rats were randomly assigned into three groups. Group one (n=12) received saline only (sham), group two (n=12) was subjected to a four-day binge (ethanol-only group), and group three (n=12) received a four-day binge with powdered Alda-1^®^ (ethanol with Alda-1^®^ group). The initial ethanol dose was 5 g/kg, and an additional dose was determined based on the following six-point intoxication behavior scale (Figure 2). For example, if the rat received a score of two, the subsequent ethanol dose was 3 g/kg (5 g minus 2 points). Food other than water was not provided during the four-day experimental period. Alda-1^®^ was administered (10 mg/kg) every 8 hours with ethanol for four days in the ethanol with Alda-1^®^ group. Powdered Alda-1^®^ was solubilized in ethanol and administered with 25% ethanol by intragastric gavage.

**Figure 2.**
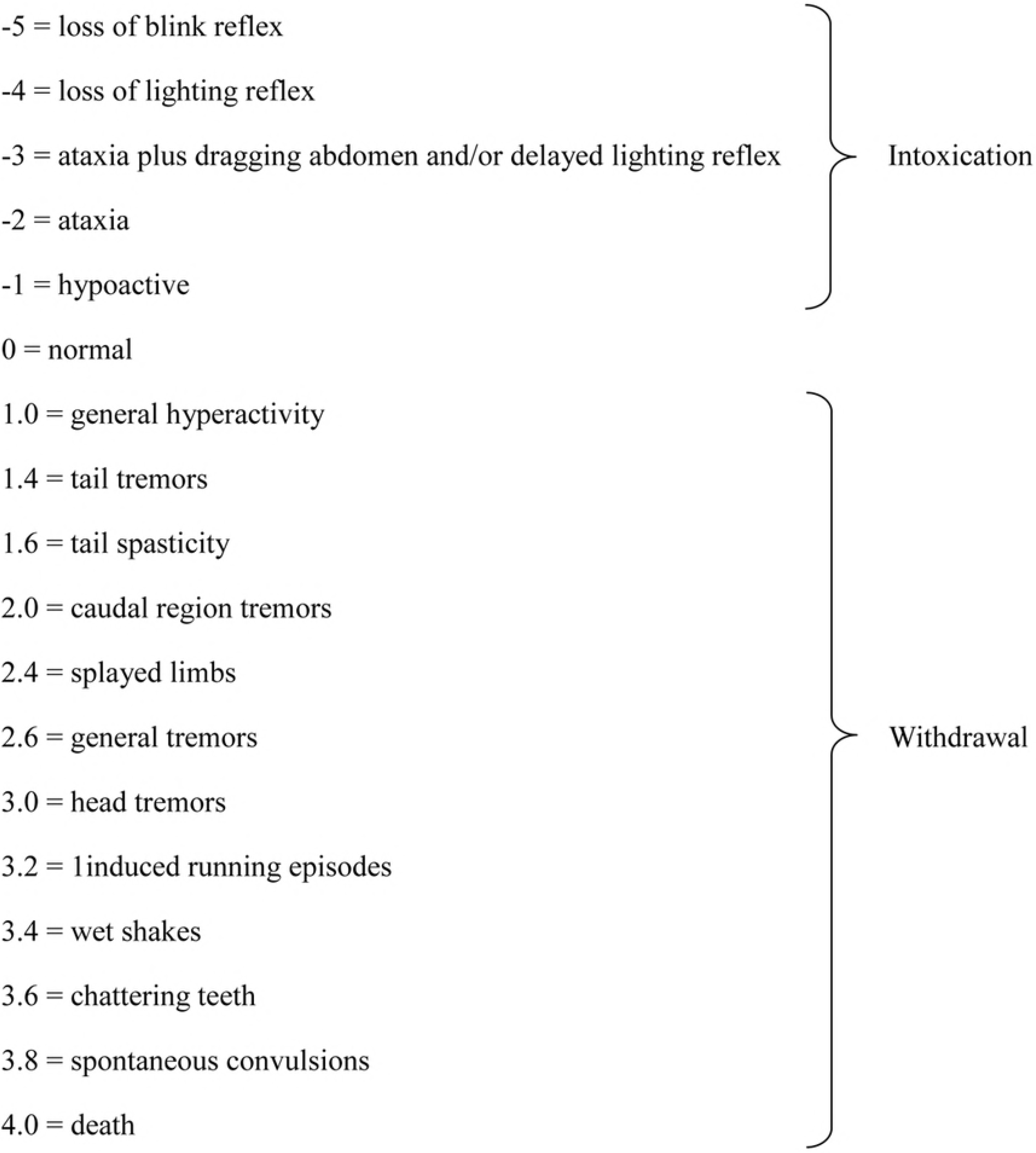
Intoxication and withdrawal behavior scores.

### Measurement of blood ALDH activity levels

We quantified ALDH enzymatic activity using Abcam’s ALDH activity assay kit (ab155893, Abcam^®^, Cambridge, United Kingdom), which detects NADH that is generated when acetaldehyde is oxidized by ALDH. NADH converts a colorless probe into a colored product that strongly absorbs light at 450 nm. An ELISA reader (iMark™ kmicroplate absorbance reader, Bio-Rad laboratories, Inc., California, USA) was used to detect the colored product with strong absorbance at 450 nm.

The blood ALDH activity of animals was measured before the first dose on the first day and 90 min after the final dose on day four (Figure 4). Eighteen animals (6 animals of each group) were randomly selected. Tail venous blood (0.3 cc) was collected and centrifuged at 3,000 RPM for 10 min.

ALDH activity was measured according to the protocol provided by Abcam ^22^.

### Behavior score

We recorded videos to observe the animals’ behavior. Videos captured for the intoxication behavior score before every ethanol injection (thrice daily) (Figure 2). Videos for the withdrawal behavior score were captured every 4 hours beginning 10 hours after the last infusion of ethanol for 72 hours. The rats’ group types were not evident in the videos. Using these videos, the blinded researcher rated the behavior (intoxication and withdrawal) scores as described by previous reports ^8, 23^ (Figure 2).

### Tissue preparation and histopathological analysis

After a withdrawal period (for three days after the last dose of ethanol), animals were anesthetized with a 2:1 ratio of zoletil (0.7 ml/kg) and rumpun (0.3 ml/kg) and were intracardially perfused with cold saline (400 ml) followed by 4% paraformaldehyde in 0.1 M phosphate-buffered saline (PBS, pH 7.4) until the upper extremities became stiff. Brains were extracted and post-fixed using the above fixative for at least four hours and were subsequently washed with tap water for 12 hours. Brains were dehydrated using ethanol and cleared using xylene. After paraffin infiltration, the brains were embedded into paraffin blocks. Six-micrometer-thick paraffin sections, including the hippocampus, were deparaffinized with xylene and dehydrated with 95% ethanol. The sections were stained with Luxol fast blue and Cresyl violet solution to detect the myelin sheath and Nissl substance using the method of Kluver et al. ^24^.

Glial fibrillary acidic protein (GFAP) and Iba-1 were immunostained in astrocytes and microglia. The deparaffinized sections were washed with 0.01 M PBS, immersed in 50% ethanol solution for 30 minutes and then washed with 0.01 M PBS again. Sections were blocked in 10% normal donkey serum (NDS, Jackson ImmunoResearch, USA) for 40 minutes and were then incubated overnight in primary antibody (GFAP: mouse, Millipore, USA, Iba-1: rabbit, Wako, Japan) diluted 1:500 in PBS. Sections were rinsed thrice in PBS and incubated for 15 minutes in 2% NDS to block the nonspecific immunoreaction of secondary antibodies. Sections that were incubated in GFAP (primary antibody) were coupled with Cy3-conjugated donkey anti-IgG (1:200, Jackson ImmunoResearch, U SA) for two hours. The Alexa Fluor 488 anti-donkey antibodies IgG (1:200 Invitrogen, USA) were used as the secondary antibody in the Iba-1 group. The samples were washed in 0.01 MPBS and distilled water (DW) and mounted using mounting medium (Vectashield, Vector Lab, USA).

### Histopathological analysis

Neuronal cell morphology as well as Iba-1-labeled microglia and GFAP-labeled astrocyte morphology was analyzed using a BX-51 Olympus Microscope and DP-70 camera (Olympus; Center Valley, PA, USA) and Visual Information Solutions image software (version 3.6.4.0, Visiopharm, Hoersholm, Denmark). Viable neurons in the Luxol fast blue and Cresyl violet-stained sections were counted at the center of the CA1 and CA2/3 fields by two blinded experienced neuropathologists. Five animals in each group were randomly selected, and the counts were determined by averaging the counts from 6 sections taken from each animal.

### Statistical analysis

Statistical analysis was performed using SPSS 18.0K software for Windows (SPSS Inc., IL, USA). All values are expressed as the mean ± SD. The p-value for statistical significance was set at 0.05. The mean value of the behavior score was compared between the ethanol-only group and the ethanol with Alda-1^®^ group using T-tests. Analysis of variance (ANOVA) was used to compare the mean value among three groups.

## Results

### General characteristics

One of twelve animals in the ethanol-only group was killed by the other animals during the intoxication period (binge). As a result, 12 animals of the sham group, 11 animals of the ethanol-only group and 12 animals of the ethanol with Alda-1^®^ group were enrolled in this study. The mean weights of animals among the three groups did not significantly differ (sham: 143.3±4.8, ethanol-only group: 145±4.0, ethanol with Alda-1^®^: 142.9±6.4 g, p=0.592).

### Behavior score

The intoxication behavioral scores (6-point scale) are presented in Figure 3, and the difference between two groups is not significant (p=0.739). The average daily dose of ethanol that was determined (adjusted) by the intoxication scores was 10.2±3.3 g/kg/day in group two (ethanol only) and 10±2.9 g/kg/day in group three (ethanol + Alda-1^®^) (Figure 3). This dose is similar to doses used in previous studies ^6, 20, 25^.

**Figure 3.**
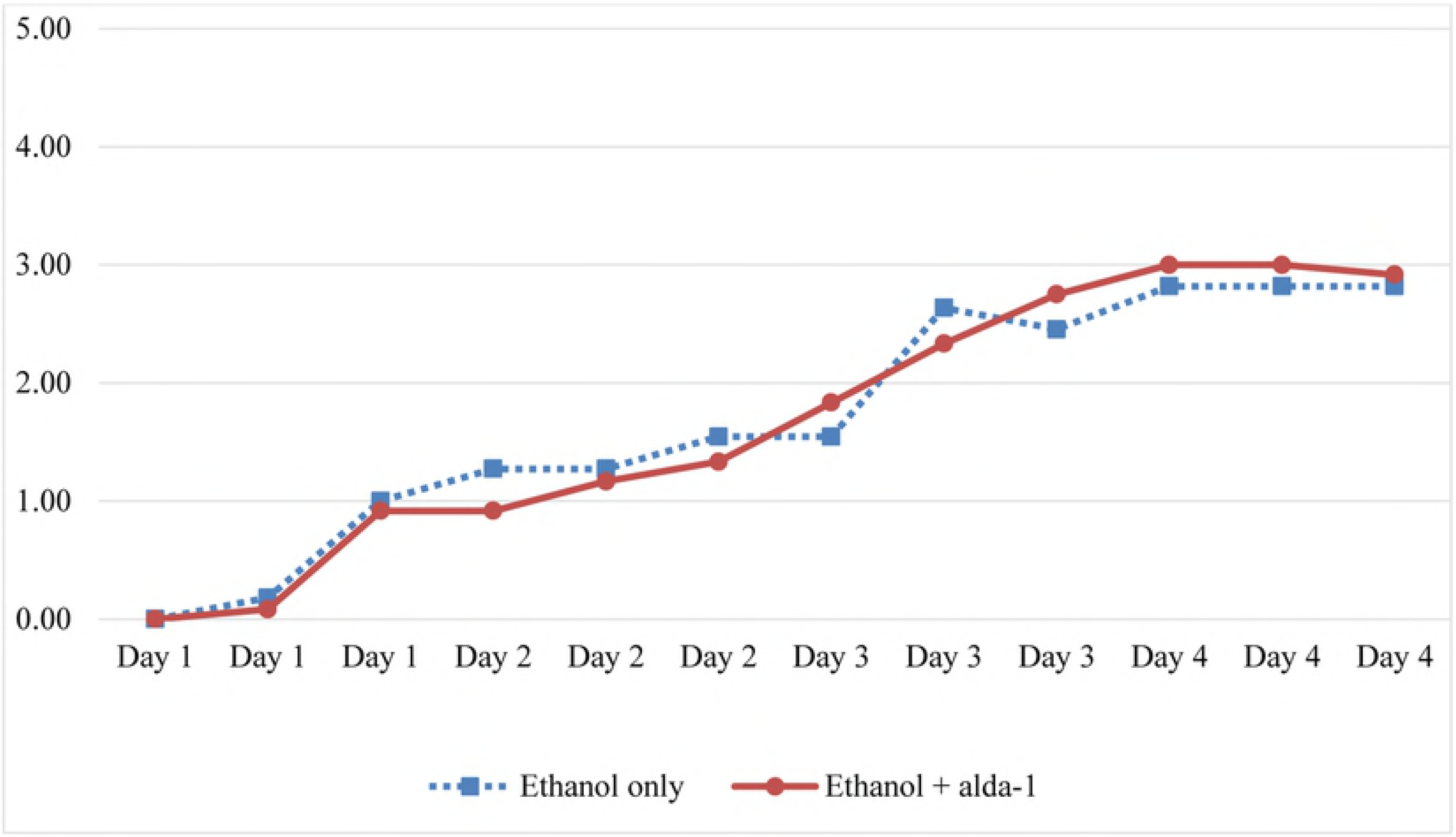
Intoxication behavior score during binge for animals in the ethanol only group and the ethanol with Alda-1^®^ group.

The withdrawal behavioral scores are presented in Figure 4. The mean withdrawal behavioral score in the ethanol-only group was significantly increased compared with the ethanol with Alda-1^®^ group, (1.56±1.30 vs. 0.76±0.90, p=<0.001). The peak for withdrawal was approximately 34 hours after the final ethanol dose. The peak withdrawal score was also increased in the ethanol-only group (3.25±0.16) compared with the ethanol with Alda-1^®^ group (2.2±0.21), and this result was significantly different (p<0.001).

**Figure 4.**
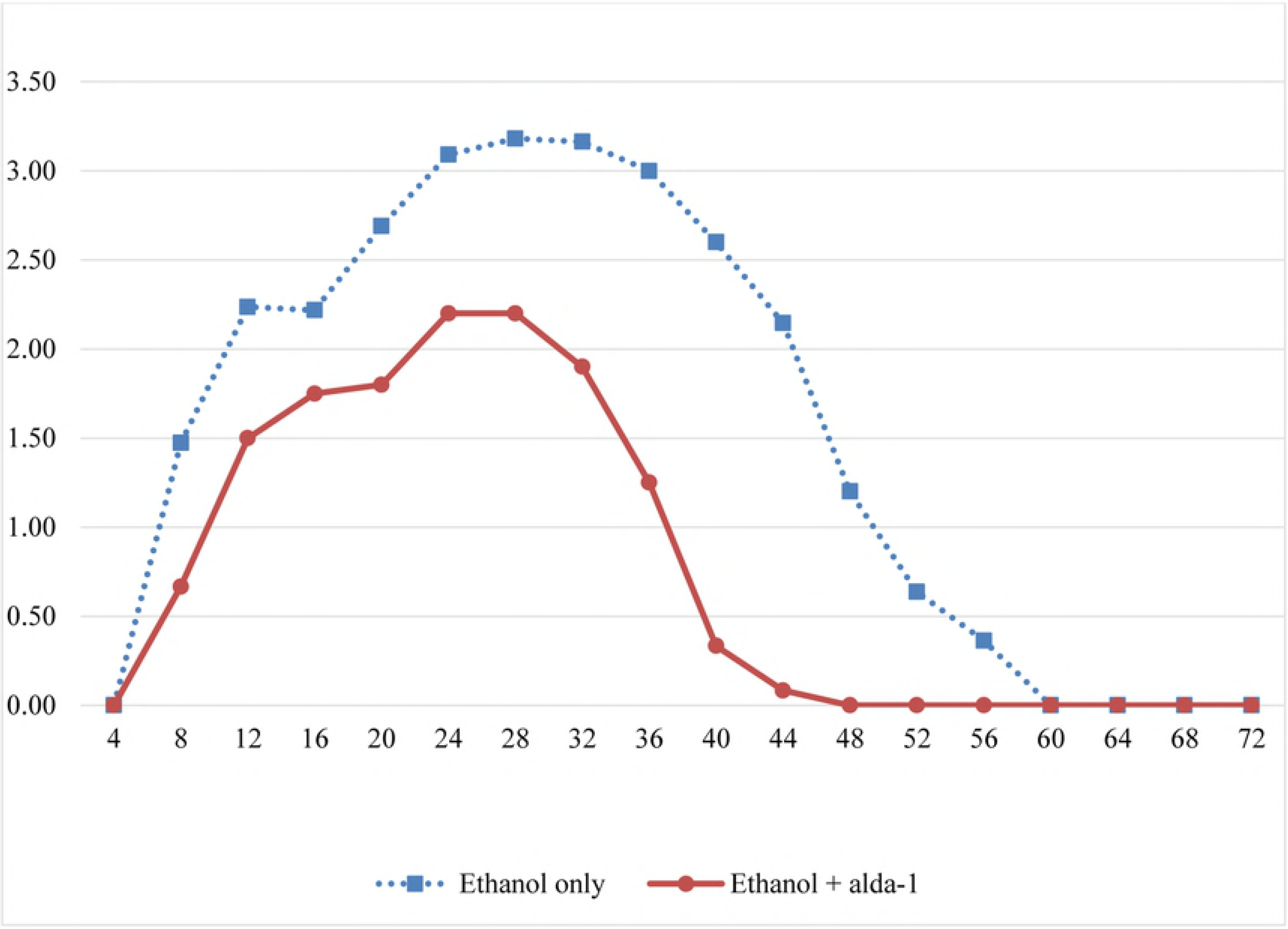
Withdrawal behavior scores in the ethanol only group and the ethanol + Alda-1^®^ group during the abstinence period.

### ALDH activity

Based on the values of the optic density measured by the enzyme-linked immunosorbent assay (ELISA) reader, and ALDH activity was calculated (Table 1). The mean baseline ALDH activities of the sham group, ethanol-only group and ethanol with Alda-1^®^ group were 4.33 (SD: 0.59), 4.23 (0.46) and 3.90 (0.57) mU/ml, respectively, and these results are not significantly different (p=0.396). After a four-day binge, the ALDH activity increased twofold in the ethanol with Alda-1^®^ group (mean: 7.87, SD: 0.67), whereas the levels of the sham group (mean: 4.07, SD: 0.53) and ethanol-only group (mean: 3.77, SD: 0.36) were slightly decreased.

**Table 1.**
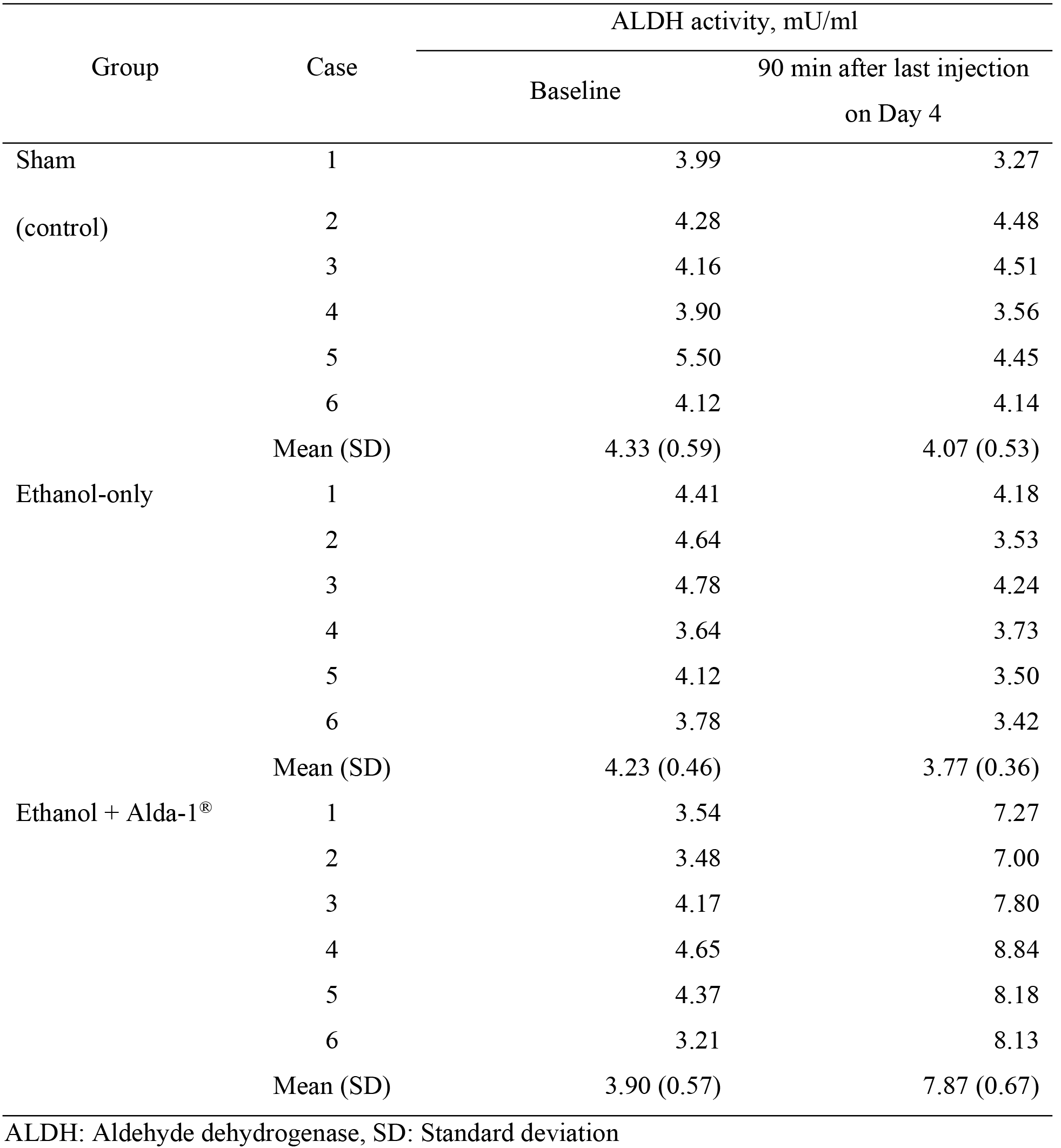
The ALDH activity among the three groups before and after binge

### Histopathology

#### Luxol fast blue and Cresyl echt violet stain

The myelinated fibers and nerve cells were dyed blue and purple, respectively. Figure 5 demonstrates distinct neuronal shrinkage, fewer neurons and loss of Nissl substance of the hippocampus, especially in the CA1 and CA2/3 fields in the ethanol-only group, whereas these features were not observed in the sham and ethanol with Alda-1^®^ group (Figure 5). Table 2 shows the mean number of viable neurons in fields from each group. The number of viable neurons in the ethanol only group was significantly decreased compared to the ethanol + Alda-1 group.

**Figure 5.**
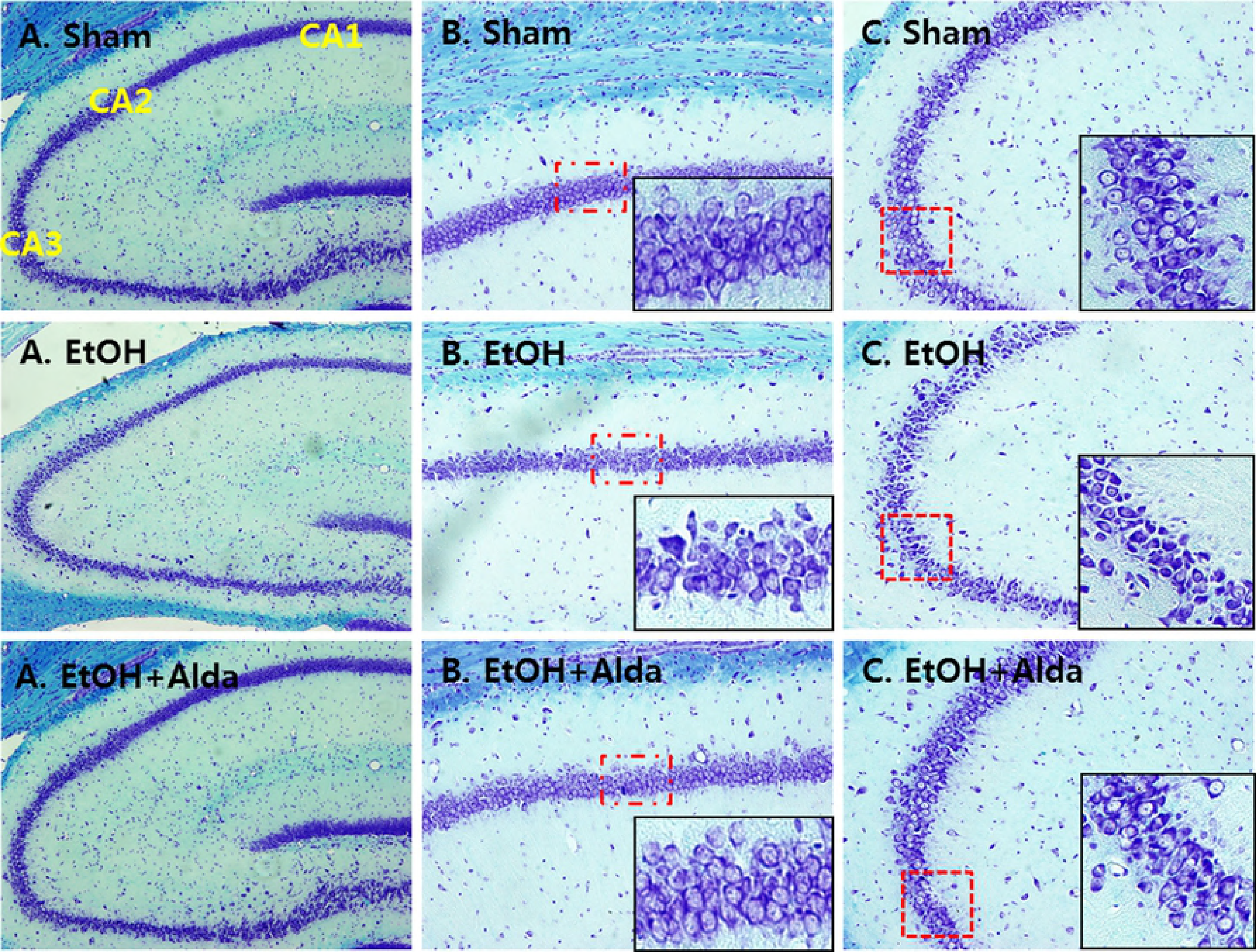
Binge ethanol exposure induces neuronal atrophy, especially in the hippocampus CA1 and CA2/3, whereas additional Alda-1^®^ administration protects against neuronal damage (Luxol fast blue and Cresyl violet stain). (A) Hippocampus (X100), (B) CA1 (X200), (C) CA2/3 (X200).

**Table 2.**
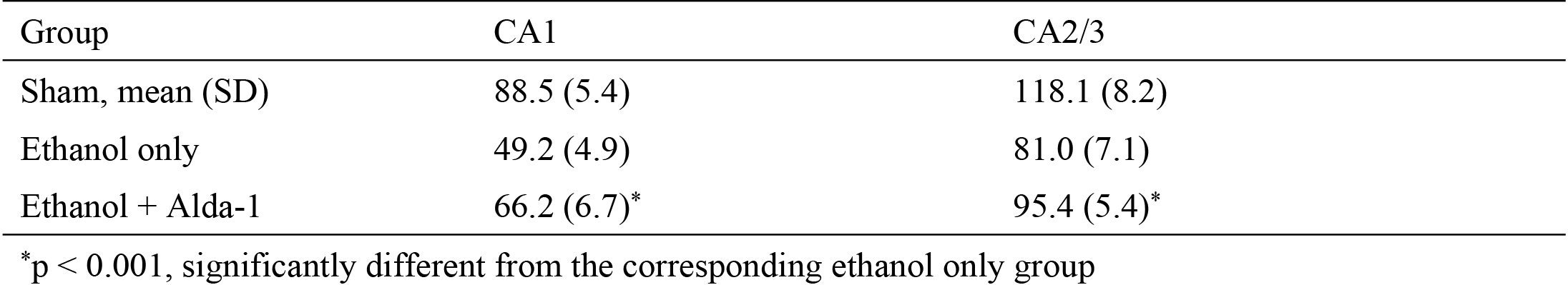
Mean number of viable neurons of the hippocampal CA1 and CA2/3 fields in each group (sham, ethanol only and ethanol with Alda-1 group)

| Group | CA1 | CA2/3 |
| --- | --- | --- |
| Sham, mean (SD) | 88.5 (5.4) | 118.1 (8.2) |
| Ethanol only | 49.2 (4.9) | 81.0 (7.1) |
| Ethanol + Alda-1 | 66.2 (6.7)* | 95.4 (5.4)* |

\*p < 0.001, significantly different from the corresponding ethanol only group

#### GFAP and Iba-1 fluorescent immunohistochemistry

The reactive expression of GFAP-a, an astrocyte marker, and Iba-1-a, a calcium binding protein expressed in microglia, was observed in field CA2/3 of the hippocampus in the ethanol-only group; however, distinct reactive astrocytosis and microgliosis were not detected in the ethanol with Alda-1^®^ group (Figure 6). Reactive astrocytosis and microgliosis are more evident in the ethanol-only group compared with the sham and ethanol + Alda-1^®^ groups.

**Figure 6.**
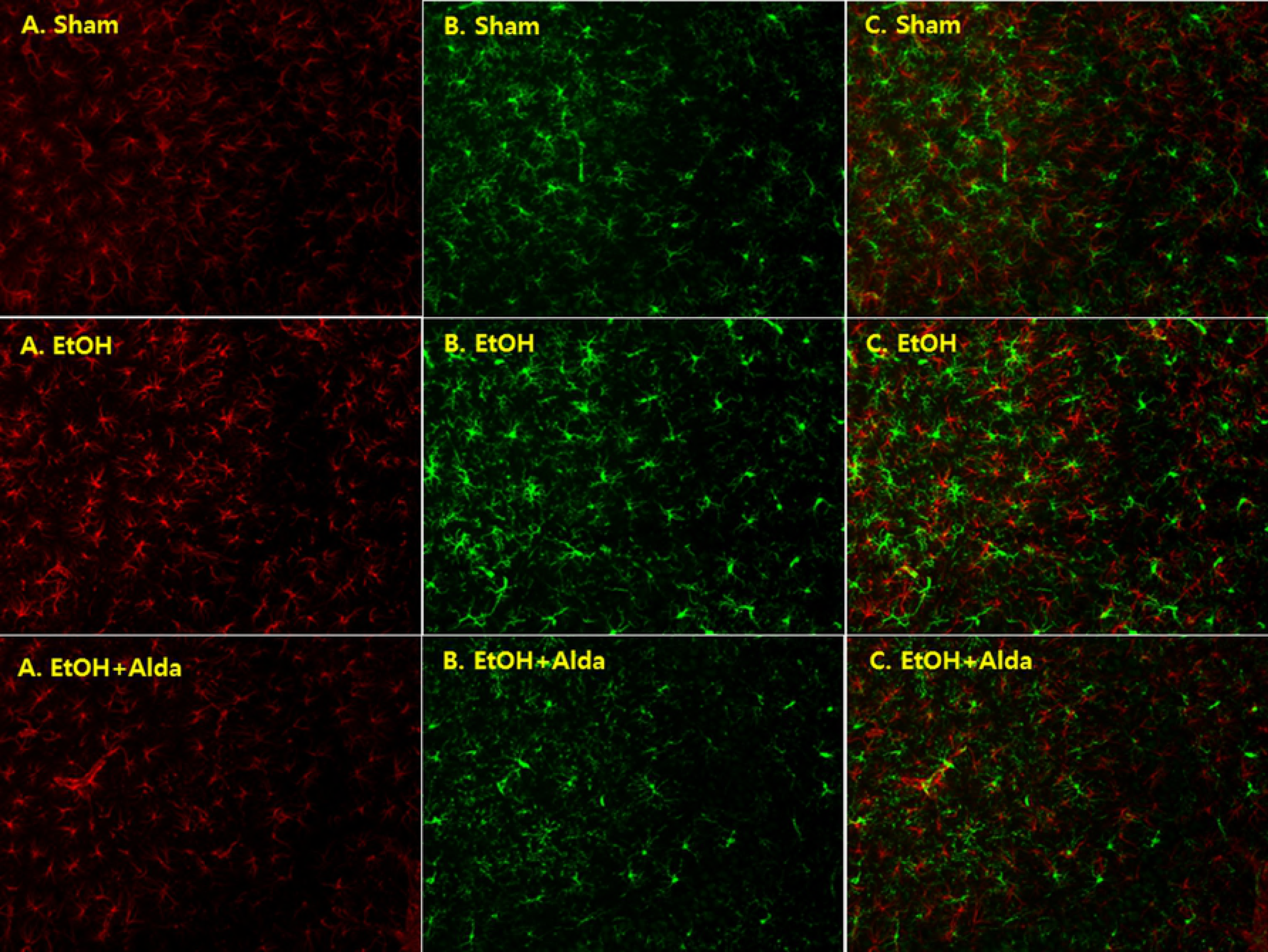
Binge ethanol exposure induces astrocyte and microglia proliferation in the hippocampus; however, these features were not distinct in the ethanol + Alda-1^®^ group. Representative images of the astrocyte marker GFAP (A) and the microglia marker Iba-1 (B) as well as the merged image (C).

## DISCUSSION

Approximately 1.9 billion adults worldwide consume at least a daily average of 13 g of ethanol (one beverage) ^26^, and approximately 140 million people consume ethanol in excess, which results in chronic alcoholism or alcohol dependence ^27, 28^. In a recent analysis, students were shown to drink heavily; 29.1% of high school students in Korea were current drinkers ^29^. Of these, 55.7% were binge drinkers, with binge drinking defined as five or more drinks per drinking episode for men and three or more drinks for women. College students in the transition period from adolescence to adulthood drink more heavily; 54.4 % of 3,371 Korean college students answered that they drank five or more drinks per one episode, and 39.7% drank once or more per week ^30^. Wechsler et al. also reported that 44% of college students binge ^31^. Adolescents are believed to be more susceptible to this binge induced-neuropathology. Several studies have demonstrated that the adolescent brain, especially the hippocampus, is more susceptible to alcohol-induced damage than the adult brain ^32, 33^.

To the best our knowledge, this is the first study to investigate the neuroprotective effect of the ALDH activator-Alda-1^®^ for binge-induced brain injury. The authors confirmed that the administration of Alda-1^®^ increases ALDH activity approximately twofold and reduces the cytotoxic damage to the brain, especially the hippocampus in adolescent rats. In this study, the binge-induced neuronal atrophy and decreased number of viable cell was confirmed via Luxol fast blue and Cresyl violet staining in the ethanol-only group, and this injury was significantly ameliorated in the Alda-1^®^-treated group, indicating that Alda-1^®^ protects against neuronal damage. These results were also supported by immunochemistry analysis of GFAP and Iba-1, which are sensitive indicators of cell injury in numerous brain injury models, including the four-day binge model ^25, 34^. Chronic alcoholism causes both cognitive dysfunction and olfactory deficits ^33, 35, 36^, and these functions are related to the above parameters; therefore the hippocampus is believed to be vulnerable to the binge-induced neuropathology ^4, 19, 20, 33^. Several studies have reported neonatal binge ethanol-induced Purkinje cell loss in the cerebellum of developing rats ^37, 38^, which depends on the developmental timing of ethanol exposure, with maximal injury achieved before postnatal day seven. However, little evidence is available in adult rat models. Binge ethanol-induced thalamic injury was also demonstrated in the developing mouse brain, but it has not been reported in adult models. Although chronic ethanol intoxication may induce cerebellar and thalamic damage ^39^, acute ethanol intoxication does not significantly impair the cerebellum and thalamus in adult rats. Thus, in this study, we selected the hippocampus to show the difference in brain injury between the binge group and binge with Alda-1 group.

As previously mentioned, ethanol-induced brain damage is largely attributed to acetaldehyde accumulation. Aldehydes are strongly electrophilic given their terminal carbonyl groups, which makes these molecules highly reactive. Aldehydes can react with cellular nucleophiles and form various adducts, which are believed to be the primary mechanism of their toxicity. Chandler et al. ^40^ suggested that glutamate-aspartate-containing pyramidal cells are potentially related to binge-induced brain damage based on the increased neuronal sensitivity to N-methyl-D-aspartate (NMDA) receptor excitation and NMDA receptor-mediated excitotoxicity in chronic alcoholism. However, several efforts to reduce binge-induced brain damage have not supported this mechanism. The NMDA antagonist MK801 does not reduce the binge-induced damage, and the glutamate AMPA receptor antagonist DNQX also does not exhibit neuroprotective effects in binge-induced damage ^41, 42^. Ke et al. ^43^ suggested that endoplasmic reticulum (ER) stress plays an important role in the pathogenesis of ethanol-induced brain damage. The ER is the site of secretory protein synthesis and folding. This organelle can respond to stress via several mechanisms (translational attenuation, upregulation of genes for ER-related proteins, and degradation of unfolded proteins). If the ER stress is prolonged or aggravated, cellular signals causing cell death are stimulated. They demonstrated that acute seven-day binge ethanol exposure significantly increased the ER stress response as indicated by the increases in activating transcription factor 6 (an ER stress-regulated transmembrane transcription factor), CCAAT-enhancer-binding proteins (C/EBP), homologous protein (CHOP, ER stress-mediated apoptosis pathway), glucose-regulated protein 78 (GRP78, a major ER chaperone), mesencephalic astrocyte-derived neutrophilic factor (MANF) and the phosphorylation of several protein kinases within 24 hours after ethanol injection. ER stress has also been implicated in neurodegenerative disorders, such as Parkinson‘s, Alzheimer’s and prion disease. Although the mechanisms of binge-induced brain damage remain unclear, it is believed that connective pathways between the frontal cortical olfactory fields and mesocorticolimbic systems are highly associated with the binge-induced brain damage.

Alda-1^®^ stimulated ALDH activity and reduced aldehyde levels in several previous studies ^17, 18^. Alda-1^®^ reduces brain injury, which is attributed to various possible mechanisms, including reduction of oxidative stress and reactive aldehydes ^16^. In this study, ALDH activity was increased twofold by Alda-1^®^. This finding is consistent with the results of Chen et al. ^17^. Although they demonstrated that Alda-1^®^ increased ALDH activity in Asian variant ALDH2 deficiency (ALDH2*2) by approximately 11-fold, we did not determine the increased ALDH level according to the genetic type because ALDH2 knock-out rats were not utilized in this study.

